# PD-1 and CTLA-4 exert additive control of effector regulatory T cells

**DOI:** 10.1101/2022.07.22.501159

**Authors:** Joseph A. Perry, Zachary Lanzar, Joseph T. Clark, Andrew P. Hart, Bonnie B. Douglas, Lindsey Shallberg, Keenan O’Dea, David A. Christian, Christopher A. Hunter

**Affiliations:** Department of Pathobiology, University of Pennsylvania, Philadelphia, PA 19104; Department of Medicine, University of Pennsylvania, Philadelphia, PA 19104

## Abstract

At homeostasis, a substantial proportion of Foxp3^+^ T regulatory cells (T_regs_) have an activated phenotype associated with enhanced TCR signals and these effector T_reg_ cells (eT_regs_) co-express elevated levels of PD-1 and CTLA-4. Short term in vivo blockade of the PD-1 or CTLA-4 pathways results in increased eT_reg_ populations, while combination blockade of both pathways had an additive effect. Mechanistically, combination blockade resulted in a reduction of suppressive phospho-SHP2 Y580 in T_reg_ cells which was associated with increased eT_reg_ proliferation, enhanced production of IL-10, and altered dendritic cell expression of CD80 and MHC-II. Thus, at homeostasis, PD-1 and CTLA-4 function additively to regulate eT_reg_ function and the ability to target these pathways in T_reg_ cells may be useful to modulate inflammation.

## Introduction

At homeostasis, Foxp3^+^ regulatory T cells (T_reg_)^1^, have a critical role in prevention of auto-immunity and can limit the intensity and duration of inflammatory responses^2–4^, yet also demonstrate heterogeneity in context of activation such as seen between central T_reg_ and effector T_reg_ cells^5,6^. T_reg_ cells can originate from the thymus (nT_reg_), or naïve CD4^+^ T cells that receive TCR stimulation combined with signals from transforming growth factor beta (TGFβ) and IL-2 can lead to Foxp3 expression and the formation of induced T_reg_ cells (iT_reg_)^7,8^. T_reg_ cells differ from conventional CD4^+^ and CD8^+^ T cells (Tconv), in that the majority of them have a TCR that recognizes self-antigens and are specialized to preserve tolerance^9–11^. It is now appreciated that T_reg_ cells require ongoing TCR activation and costimulation to retain Foxp3 expression, suppressive capacity^12^, and survival^5,13,14^. This is illustrated by the spontaneous immunopathology in experimental models when T_reg_ cells are absent^15–18^. The clinical relevance of T_reg_ cell mediated control of adaptive responses is illustrated by X-linked immunodysregulation polyendocrinopathy and enteropathy (IPEX). In these patients, the Foxp3 gene is mutated and has an impaired ability to drive T_reg_ formation, resulting in autoimmune diseases such as neonatal type 1 diabetes, hemolytic anemia, eosinophilia, and hyper IgE production^19^.

Given the role of T_reg_ cells in limiting immune responses there is considerable interest in promoting their activities to limit inflammation while the ability to antagonize T_reg_ cells is one approach to augment anti-tumor responses^20–22^. There is emerging evidence that the relative ratio of effector Tconv cells: T_reg_ cells is an important determinant for the outcome of immunotherapy in cancer^23^. However, too many T_reg_ cells can be deleterious and lead to reduced effector responses in the context of infection or cancer^22,24^. Consequently, there need to be mechanisms to balance T_reg_ cell activities such as IL-2 availability as a mechanism involved in modulation of the T_reg_ cell pool^25,26^. In addition, there is evidence that the inhibitory receptors PD-1^27,28^ and CTLA-4^21^ restrict T_reg_ cell activities in the setting of cancer, autoimmunity and infection^21,22,28^.

PD-1 and CTLA-4 are expressed by activated T cells and most studies on these pathways have focused on their impact on effector responses which has formed the basis for checkpoint blockade in cancer. In this context, there is evidence of PD-1 and CTLA-4 to act in cis and engage SHP2 phosphatases^29–31^ antagonize TCR signals^32–34^, and thus blunt the response of effector T cells^35,36^. In addition, the ability of the extracellular domain of CTLA-4 to sequester CD80/86 provides an additional trans mechanism to limit accessory cell function required for optimal effector T cell activities^32^. A subset of T_reg_ cells also express these receptors^22,28^, and several reports have highlighted that effector (eT_reg_) cells express the highest levels of PD-1 and CTLA-4^27,28^. It appears that eT_reg_ cells receive continuous TCR signals, but constitutive signals through PD-1 constrain the size of the eT_reg_ cell pool^27,28^. These studies raise questions about the relationship between the PD-1 and CTLA-4 pathways and whether mitigation of these checkpoint proteins impact the ratios of cT_reg_ : eT_reg_ cell populations. The studies presented here reveal at homeostasis that the combined blockade of PD-1 and CTLA-4 have an additive effect on expansion of eT_reg_ cell populations associated with reduced accessory cell function. Thus, PD-1 and CTLA-4 have distinct but complementary roles in the tonic regulation of T_reg_ cell homeostasis.

## Materials and methods

### Mice

All mice used were housed in the University of Pennsylvania Department of Pathobiology vivarium with 12 hour light and dark cycles, maintained at temperature ranges of 68°F - 77°F and humidity ranges from 35% - 55% humidity in accordance with institutional guidelines.

C57BL/6 mice were purchased from Taconic (Rensselaer, NY, USA) at 6 weeks of age and housed in the University of Pennsylvania Department of Pathobiology vivarium for 2 – 4 weeks until used.

*Ethical oversight of all animal use in this study was approved by the University of Pennsylvania Institutional Animal Care and Use Committee*.

### In vivo combination checkpoint blockade

#### In vivo blockade antibodies

Details of antibodies and reagents in blockade can be found in **Supplementary Table 1**.

Inhibition of PD-1/PD-L1 signaling was performed by intraperitoneal injection of 1mg/dose of αPD-L1 (clone: 10F.9G2, BioXcell) supplemented with 500μg/dose of polyclonal hamster IgG isotype (clone: polyclonal Armenian hamster, BioXcell). Inhibition of CTLA-4 signaling was performed by intraperitoneal injection of 500μg/dose of αCTLA-4 (clone: UC10-4F10-11, BioXcell) supplemented with 1mg/dose of IgG2b isotype (clone: LTF-2, BioXcell) while control mice were treated with 1mg/dose IgG2b isotype supplemented with 500μg/dose of polyclonal hamster IgG isotype. Mice were killed 72 hours following treatment and splenocytes were analyzed via flow cytometry.

### Tacrolimus Treatment

FK506 (F4679-5MG, Sigma-Aldrich, MO, USA) was reconstituted in DMSO to 25mg/ml, and then the reconstituted stock was diluted in 1xDPBS to achieve a working concentration of 2.5mg/ml. 8-week-old C57BL/6 mice were subcutaneously injected with 50µl of FK506 at 2.5mg/ml to deliver 125µg of FK506 per dose daily of either FK506 or PBS vehicle control every 24 hours over a 96 hour period. Following 96 hours of treatment, splenocytes were then harvested and analyzed via flow cytometry.

### Isolation of tissues for analysis

#### Tissue Preparation

Single cell suspensions were prepared from spleen for flow cytometry analysis. Spleens were mechanically processed and passed through a 70µm nylon filter and then lysed in 1ml of 0.846% solution of NH4Cl for red blood cell lysis. The cells were then washed in cRPMI and stored on ice.

### Analysis by flow cytometry

#### Staining antibodies and staining reagents

Antibody, viability dye, Fc block, dilutions, and buffer reagent details can be found on **Supplemental Table 1**.

#### T cell staining

Aliquots consisting of 5e6 cells were washed with ice cold 1xDPBS in a 96 well round bottom plate, then incubated in in 50µl volume of viability stain reconstituted in 1xDPBS for 20 minutes on ice and then washed in 0.2% FACS buffer. The cells were then incubated in 50µl volume of Fc block for 30 minutes on ice. The cells were washed in 0.2% FACS buffer, and then incubated for 30 minutes on ice in 50µl volume of antibody cocktail composed of surface-stain antibodies in 0.2% FACS buffer supplemented with brilliant stain buffer **(Supplemental Table 1)**.

The cells were washed in 0.2% FACS buffer and re-suspended in 100µl Foxp3 Perm-fix cocktail (00-5523-00, Thermo Fisher Scientific) for 4 hours at 4°C. The cells were then washed twice in 1X permeabilization buffer, and then re-suspended in an intracellular staining cocktail composed of intracellular-stain antibodies diluted in 1x permeabilization buffer supplemented with normal goat serum of for 2 hours at 4°C. The cells were then washed with 1x permeabilization buffer twice, and then resuspended in 50µl of Goat α-Rabbit detection antibody diluted in 1X permeabilization buffer for 2 hours at 4°C. The cells were washed in 1x permeabilization buffer and resuspended in 500µl 0.2% FACS buffer for flow cytometric analysis.

#### Cytokine staining

To detect intracellular cytokines on T cells, cells were re-suspended in a 1X dilution of Cell Stimulation Cocktail Plus Protein Transport Inhibitors (Invitrogen, #00-4975-93, CA) in cRPMI for 2 hours at 37°C and 5% CO_2_. Cells were then washed, surface stained, and permeabilized as described above in the T cell panel. The cytokine stain prepped cells were then intracellularly stained with a cytokine detection panel for 2 hours on ice. The cells were washed and then resuspended in 500µl 0.2% FACS buffer for analysis.

#### Myeloid staining

Aliquots of 5e6 cells were washed in ice cold 0.2% FACS buffer in a 96 well and then viability stained and Fc-blocked as described in the T cell panel. The cells were surface stained in 50µl of antibody cocktail consisting diluted in 0.2% FACS buffer supplemented with brilliant stain buffer on ice for 30 minutes. The cells were washed and fixed in with 2% PFA (15710-S, Electron Microscopy Sciences) diluted in 0.2% FACS buffer for 15 minutes at room temperature. The cells were then washed and then re-suspended in 500µl 0.2% FACS buffer for analysis.

#### Phos-flow

Splenocyte-derived CD4^+^ T cells were isolated using Easysep Mouse CD4^+^ T cell isolation kit (19852, STEMCELL Technologies), and then 2e5 cells/well were plated in a 96 well plate, and viability stained as described above using sterile 1xDPBS. Cells were blocked for PD-1, CTLA-4, or combination of PD-1 and CTLA-4 using anti-PD-1 (clone: RMP1-14, BioXcell), anti-CTLA-4 (clone: UC10-4F10-11, BioXcell) or isotype control antibodies (clone: 2A3, BioXcell, and clone: polyclonal Armenian hamster IgG, BioXcell). The cells were blocked in 100µl of PD-1/CTLA-4 blocking cocktails in sterile MACS buffer (2% FCS, 2mM EDTA, in 1xDPBS) at a concentration of 10µg/ml of antibody on ice for 20 minutes. The cells were washed with sterile MACS and were then resuspended in 100µl sterile RPMI containing 0.5% BSA, and then transferred to a 96 well plate that had been coated overnight at 4°C with 5µg/ml αCD3 (BE0001-1, BioXcell), 5µg/ml CD80-Fc (555404, Biolegend), and 2µg/ml PD-L1-Fc (758206, Biolegend). The cells were either incubated at 37°C for 30 minutes or one hour, and then mixed with 100µl of 5% PFA (15710-S, Electron Microscopy Sciences) diluted in ice cold 1xDPBS and incubated on ice for 20 minutes (direct exvivo phos-flow assessments were directly fixed without incubation). The cells were washed 2x in 1xDPBS, and permeabilized in 100µl Foxp3 Perm-fix cocktail (00-5523-00, Thermo Fisher Scientific) for 2 hours, and then washed as described above. The cells were re-suspended in an intracellular staining cocktail composed of intracellular-stain antibodies diluted in 1x permeabilization buffer for 2 hours at 4°C. The cells were washed twice in 1x permeabilization buffer and resuspended in 0.2% FACS buffer for flow cytometric analysis.

#### Data acquisition

The cells were analyzed on a FACS Symphony A5 (BD Biosciences) using BD FACSDiva v9.0 (BD Biosciences) and analysis was performed with FlowJo (10.8.1, BD biosciences).

#### Statistics

Statistical analysis was performed using Prism 9 for Windows (version 9.2.0). For comparison of means between two groups, either a two-tailed unpaired, or paired student’s *t* test was utilized with a 95% CI depending on separate treatment groups or treatments within groups. Analysis for univariate statistics comparing multiple means was performed using a one-way ANOVA (family-wise significance and confidence level of 95% CI), with post-hoc analysis consisting of Fisher’s LSD test for direct comparison of two means within the ANOVA, or Tukey’s multiple comparisons test for comparisons of all means within the test group for multiple-comparison correction. For multi-group multivariate analysis, a two-way ANOVA with post-hoc analysis utilizing Sidak’s multiple comparisons test for comparisons across two groups with two variables, or Tukey’s multiple comparisons test for comparisons across multiple groups for multiple variables (also with a 95% CI). Probability for *p* values <0.05 or lower were considered statistically significant. All error bars in the figures indicate standard error of the mean (SEM).

#### UMAP analysis

Uniform Manifold Approximation and Projection for Dimension Reduction (UMAP) analysis was performed using the UMAP plug-in using the Euclidean distance function with a nearest neighbor score of 20, and a minimum distance rating of 0.5 (version: 1802.03426, 2018, ©2017, Leland McInness) for Flowjo (Version 10.8.1). All stained parameters were included in UMAP analysis except for: Live Dead (gated out), CD4 (pre-gated), PD-L1 and CTLA-4 (avoiding grouping bias), Foxp3 (avoiding grouping bias or already pre-gated). The heatmap overlay figures for UMAP analysis presented are based on median fluorescence of each labeled stain in each figure and generated within Flowjo (Version 10.8.1).

#### Data availability statement

The data that support the findings of this study are available on request from the corresponding author C.A. Hunter.

## Results

### Preferential expression of PD-1 and CTLA-4 by eT_reg_ cells

To compare the relative activation state of CD8^+^ T cells, CD4^+^ Foxp3^-^ T cells (Tconv) and CD4^+^ Foxp3^+^ cells (T_reg_s) at homeostasis, the levels of CD69, CD11a, and CD44 (markers associated with TCR activation) were assessed. T_reg_ cells had highest expression of CD69, CD11a, and CD44 **(Figure 1A, Supplemental Figure 1A)**, and the highest proportion of CD11a^hi^ CD44^hi^ cells **(Figure 1B)**. Likewise, T_reg_ cells also had the largest proportion of Ki67^+^ and cMyc^+^ cells, two markers associated with proliferation^38,39^ **(Figure 1C)**. These markers of activation and proliferation correlated with the preferential co-expression of PD-1 and CTLA-4 by T_reg_ cells compared to non-T_reg_ T cells **(Figure 1D)**. Next, T_reg_ cells were divided into PD-1^-^ CTLA-4^low^ and PD-1^+^ CTLA-4^hi^ T_reg_ cells **(Supplemental Figure 1B)**, that correlate with central (cT_reg_) and effector (eT_reg_) subsets^5,28^ respectively compared with the PD-1^-^ CTLA-4^low^ cT_reg_ cells, the PD-1^+^ CTLA-4^hi^ eT_reg_ cells had significantly greater expression of CD69, CD11a, CD44, and Helios **(Figure 2A)**. As expected, the PD-1^+^ CTLA-4^+^ c eT_reg_ subset was enriched for cells that co-expressed elevated levels of CD11a and CD44 **(Figure 2B)**, Ki67 and cMyc **(Figure 2C)**, and this subset had an increased ability to produce IL-10 **(Figure 2D)**. We also noted that the proportion of these proliferative eT_reg_ cells increased with age and could be as high as 40% of the T_reg_ cells in older mice **(Supplemental Figure 1C)**. Moreover, when eT_reg_ and cT_reg_s were stained for phosphorylation of TCR-associated ZAP70, PI3k, AKT, ERK1/2, and mTOR, in addition to both SHP2 tyrosine sites Y542 and Y580, (of which Y542 can dephosphorylate Y580 - the active tyrosine site associated with inhibition of TCR signals^35,40,41^), eT_reg_ cells had the highest levels of TCR-associated and SHP2 phospho-proteins **(Figure 2E, F)**. These results suggest that eT_reg_ cells receive constitutive TCR activation while experiencing ongoing SHP2 mediated restriction of these signals.

**Figure 1:**
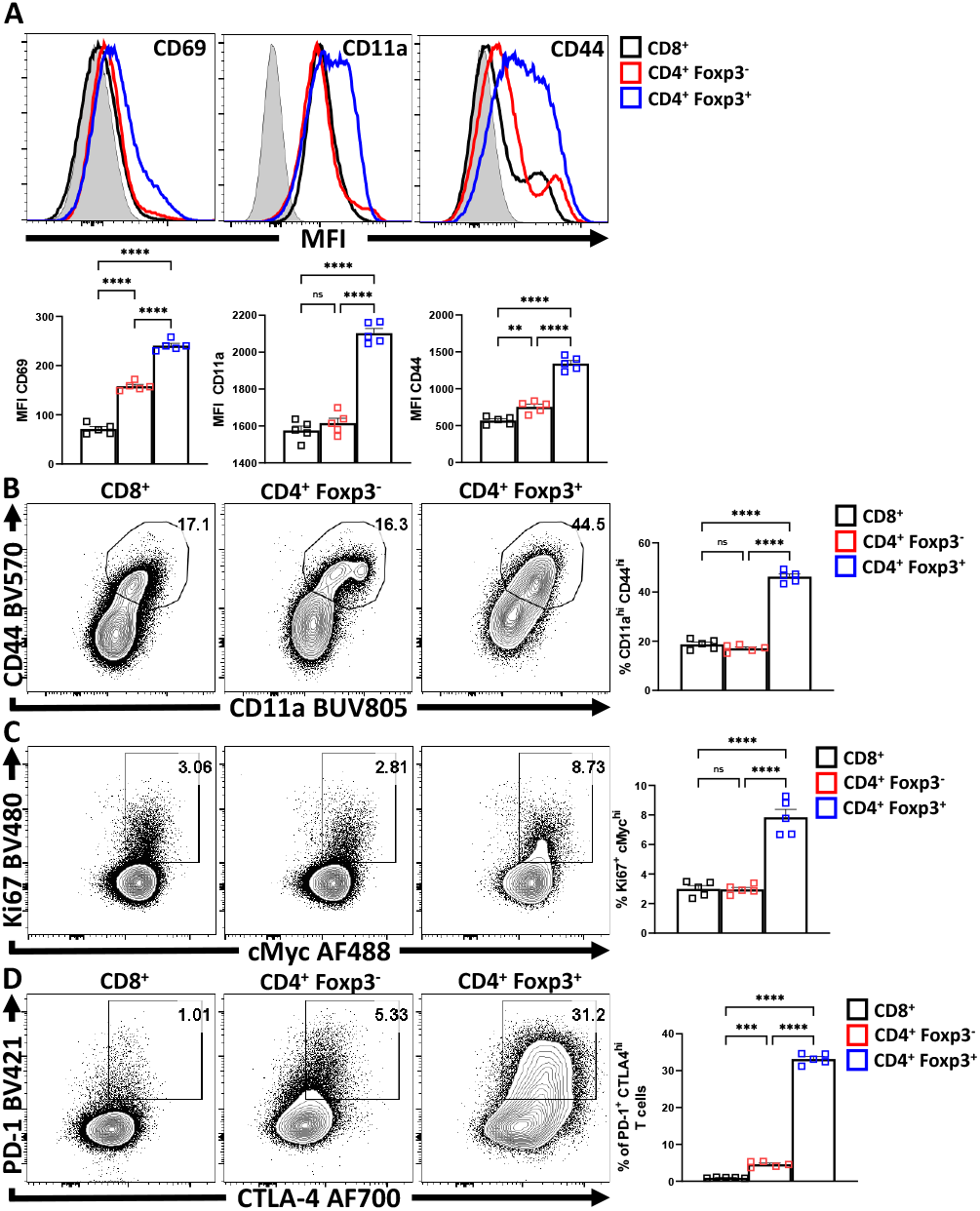
T_reg_ cells are the most active and proliferative T cells at homeostasis, yet express PD-1 and CTLA-4. Splenocytes from naïve 8-week old male C57BL/6 mice were analyzed via high-parameter flow cytometry to compare the expression of activation, proliferation, and PD-1/CTLA-4 proteins CD8^+^, CD4^+^ Foxp3^-^ (CD4 Tconv), and CD4^+^ Foxp3^+^ (T_reg_) cells for the following figures. **(A)** Histogram comparisons of gMFI of CD69, CD11a, and CD44 expression between the CD8/CD4^+^ Tconv and T_reg_ compartments *(n = 5/group, 1-way ANOVA with Tukey’s multiple comparisons test, ** = p < 0*.*01, **** = p < 0*.*0001, 4 experimental replicates)*. **(B)** Flow plots of ex-vivo CD11a and CD44 staining comparing the proportion of CD11a^hi^ CD44^hi^ cells within each subset *(n = 5/group, 1-way ANOVA with Tukey’s multiple comparisons test, **** = p < 0*.*001, 4 experimental replicates)*. **(C)** Plots of depicting comparisons of the proportion of Ki67^+^ cMyc^hi^ cells across these subsets *(n = 5/group, 1-way ANOVA with Tukey’s multiple comparisons test, **** = p < 0*.*0001, 4 experimental replicates)*. **(D)** Plots demonstrating proportions of PD-1^+^ and CTLA-4^hi^ cells between the Tconv and T_reg_ compartments *(n = 5/group, 1-way ANOVA with Tukey’s multiple comparisons test, *** = p < 0*.*001, **** = p < 0*.*0001, 4 experimental replicates). All data presented are means +/- SEM and show individual data points*.

**Figure 2:**
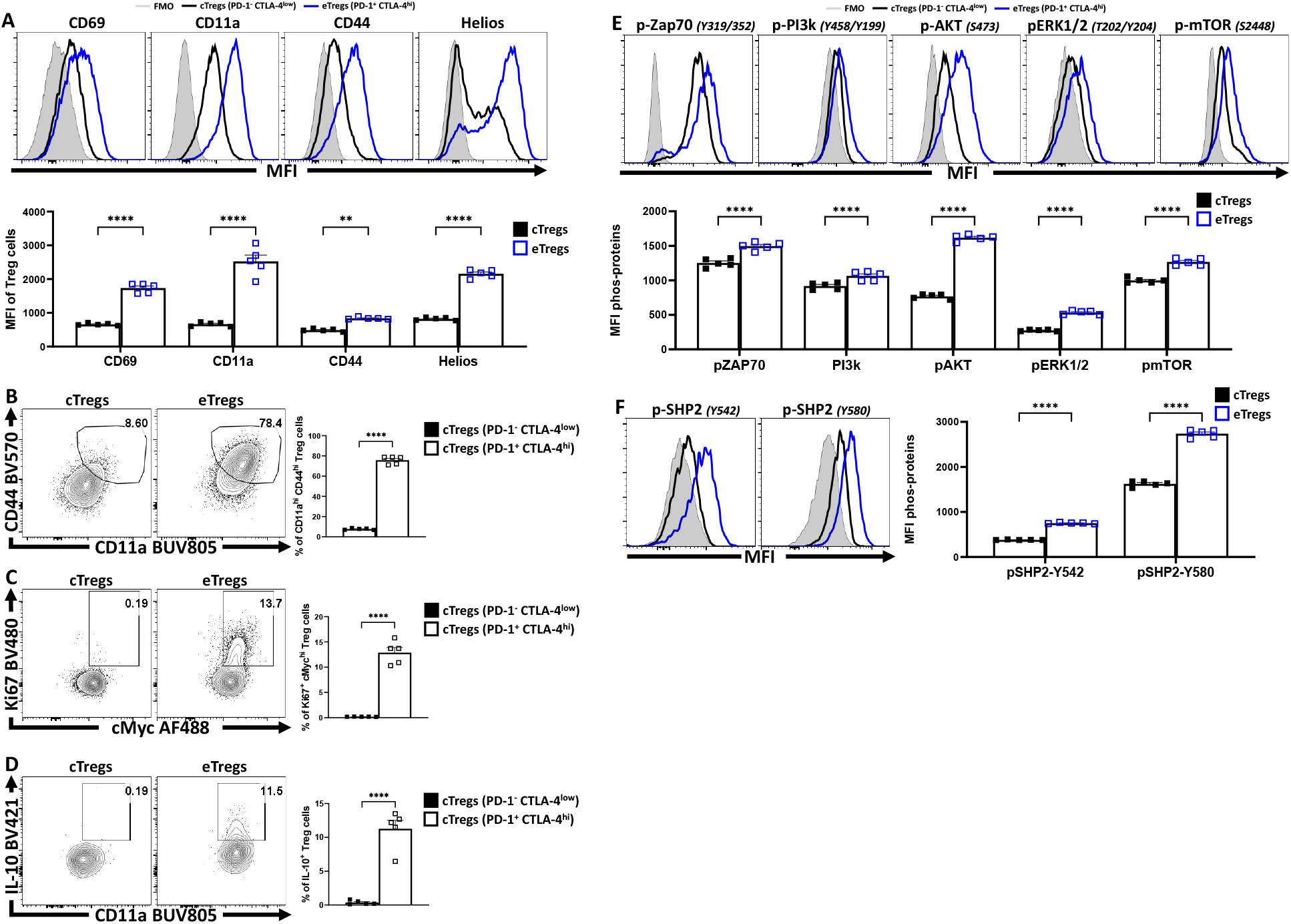
PD-1^+^ CTLA-4^+^ eT_reg_ cells express Helios and have activated T_reg_ effector phenotypes compared to PD-1^-^ CTLA-4^-^ cT_reg_ cells. Splenocytes from naïve 8-week old male C57BL/6 mice were analyzed via high-parameter flow cytometry and then pre-gated **(Supplemental Figure 1B)** on PD-1^+^ CTLA-4^hi^ (eT_reg_) vs PD-1^-^ CTLA-4^low^ (cT_reg_) subsets. Phenotypes were compared between the c/eT_reg_ subsets based on the expression of proteins associated with activation, proliferation, and IL-10 production. Additionally, TCR-downstream phosphorylation potential in response to activation between T_reg_ subsets was also evaluated. **(A)** Comparative histograms of CD69, CD11a, CD44, and Helios between eT_reg_ and cT_reg_ subsets demonstrating greater expression of activation associated proteins and Helios on eT_reg_ cells *(n = 5/group, 2-way ANOVA with Sidak’s multiple comparisons test, ** = p < 0*.*01, **** = p < 0*.*0001, 4 experimental replicates)*. **(B)** Ex-vivo flow-plots comparing the proportion of CD44^hi^ CD11a^hi^ populations and Ki67^+^ cMyc^hi^ populations **(C)**, between cT_reg_ and eT_reg_ subsets *(n = 5/group, two-tailed unpaired student’s t-test, **** = p < 0*.*0001 4 experimental replicates)*. **(D)** Flow plots following PMA/Ionomycin stim comparing the proportion of IL-10^+^ CD11a^hi^ cells between cT_reg_ and eT_reg_ subsets *(n = 5/group, two-tailed unpaired student’s t-test, **** = p < 0*.*0001 4 experimental replicates)*. **(E)** Histogram comparisons of gMFI of p-ZAP70, p-AKT, pERK1/2, and p-mTOR of cT_reg_ and eT_reg_ cells exvivo,demonstrating a greater magnitude of phospho-protein presence in eT_reg_ cells comparatively *(n = 5/group, 2-way ANOVA with Sidak’s multiple comparisons test, **** = p < 0*.*0001, 2 experimental replicates)*. **(F)** Histogram comparisons of gMFI of p-SHP2 for Y580 and Y542 residues exvivo on cT_reg_ and eT_reg_ cells *(n = 5/group, 2-way ANOVA with Sidak’s multiple comparisons test, **** = p < 0*.*0001, 2 experimental replicates). All data presented are means +/- SEM and show individual data points*.

### Homeostatic blockade of PD-L1 and CTLA-4 enhances the eT_reg_ compartment

Previous studies showed that blockade of PD-L1 at homeostasis enhanced T_reg_ cell responses^28^. To determine whether CTLA-4 also plays a similar role and how it relates to PD-1, cohorts of 8-week-old C57BL/6 mice were treated with a single intraperitoneal injection of control antibodies alone or in combination with α-PD-L1, α-CTLA-4, or a combination of α-PD-L1 and α-CTLA-4. Splenocytes from these hosts were harvested 72 hours later and analyzed via flow cytometry. The blockade of PD-L1 or CTLA-4 resulted in a significant enrichment in the proportion and total number of T_reg_ cells, yet when these blocking antibodies were combined there was an additive increase in the number of T_reg_ cells **(Figure 3A)**. This was accompanied by a concurrent increase in the proportion and total number of activated (CD11a^hi^ CD44^hi^) eT_reg_-associated cells, which correlated with the observed total increase in T_reg_ cells **(Figure 3B)**. This short-term blockade of the PD-1 and CTLA-4 pathways did not impact the non-T_reg_ subsets (CD4^+^ Tconv, and CD8^+^ T cells) but resulted in increases in activated (CD11a^hi^ CD44^hi^) T_reg_ cells with further increases in the co-blockade treated hosts **(Figure 3B)**. The enrichment of activated eT_reg_ cells correlated with increases in PD-1^+^ CTLA-4^hi^ T_reg_ cells with either blockade and when PD-1 and CTLA-4 were simultaneously blocked there was an additive increase in the ratio of eT_reg_ cells to cT_reg_ cells **(Figure 3C)**.

**Figure 3:**
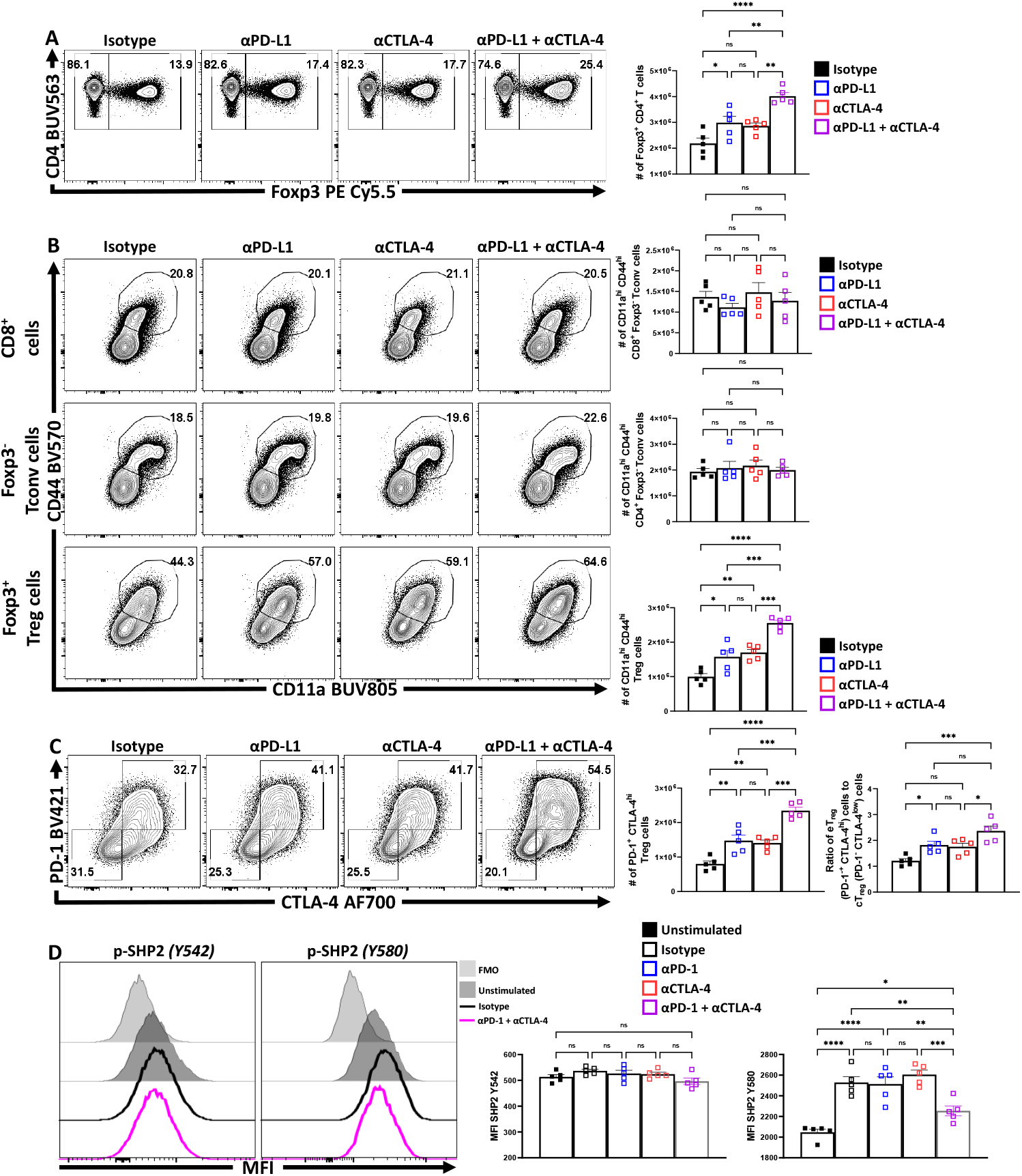
PD-1 and CTLA-4 additively restrict activated eT_reg_ cells at homeostasis. Cohorts of 8 week-old male C57BL/6 mice were given a single intraperitoneal injection of either αPD-L1, or αCTLA-4, or combination αPD-L1 and αCTLA-4, or Isotype control antibody. Splenocytes were harvested for analysis 72 hours later and analyzed via high parameter flow-cytometry. **(A)** Flow plots of bulk CD4^+^ T cells demonstrating increases in the proportion and number of T_reg_ cells following either blockade, with greatest enrichments occurring with combination blockade *(n = 5/group, 1-way ANOVA with Tukey’s multiple comparisons test, * = p < 0*.*05, ** = p < 0*.*01, *** = p < 0*.*001, **** = p < 0*.*0001, 3 experimental replicates)*. **(B)** Comparison of the proportions and number of CD44^hi^ CD11a^hi^ populations between CD8^+^ Tconv, CD4^+^ Tconv, and T_reg_ cells following 72 hours of single or combination α-PD-L1/CTLA-4 checkpoint blockade treatment *(n = 5/group, 1-way ANOVA with Tukey’s multiple comparisons test, ** = p < 0*.*01, *** = p < 0*.*001, **** = p < 0*.*0001, 3 experimental replicates)*. **(C)** Flow plots of cT_reg_ (PD-1^-^ CTLA-4^low^), and eT_reg_ (PD-1^+^ CTLA-4^hi^) cells following blockade treatment, demonstrating enrichment of PD-1^+^ CTLA-4^hi^ cells with either blockade, with the greatest enrichment occurring when both pathways were blocked *(n = 5/group, 1-way ANOVA with Tukey’s multiple comparisons test, ** = p < 0*.*01, *** = p < 0*.*001, **** = p < 0*.*0001, 3 experimental replicates)*, and subsequent ratio of eT_reg_ to cT_reg_ cells that were shifted with treatment *(n = 5/group, 1-way ANOVA with Tukey’s multiple comparisons test, * = p< 0*.*05, ** = p < 0*.*01, *** = p < 0*.*001, **** = p < 0*.*0001, 3 experimental replicates)*. **(D)** Enriched bulk CD4^+^ T cells were treated with either αPD-1, or αCTLA-4, or combination αPD-1 and αCTLA-4, or Isotype control antibody, and then stimulated with plate-bound α-CD3, PD-L1-Fc, and CD80-Fc and phospho-stained. Depicted are histogram comparisons of the T_reg_ subset (CD4^+^ Foxp3^+^) comparing gMFI of p-SHP2 at tyrosine residues Y542 and Y580 on T_reg_ cells *(n = 5/group, 1-way ANOVA with Fisher’s LSD individual comparisons test, * = p < 0*.*05, ** = p < 0*.*01, *** = p < 0*.*001, **** = p < 0*.*0001, 2 experimental replicates). All data presented are means +/- SEM and show individual data points*.

Next, the impact of combination blockade on phosphorylation of suppressive SHP2 tyrosine phosphatases during activation was considered. SHP2 tyrosine phosphatase activity restricts CD28-mediated co-stimulation^40^, and there are two tail tyrosine residues; Y542, which mitigates SHP2 phosphatase activity and Y580, which stimulates suppressive SHP2 phosphatase activity ^41^. To evaluate the impact of PD-1 and CTLA-4 blockade on SHP2 phosphorylation splenocyte-derived MACS enriched CD4^+^ T cells **(Supplemental Figure 2A)** were first treated with either an isotype control, α-PD-1, α-CTLA-4, or α-PD-1 plus α-CTLA-4. These cells were transferred to plates coated with PD-L1-Fc, CD80-Fc, and α-CD3 in serum-free media. After incubating the cells for 1 hour, the cells were then fixed and phosphorylation of SHP2 tyrosine residues Y542 and Y580 were measured via flow cytometry, subsetting on Foxp3^+^ T_reg_ cells. Firstly, T_reg_ cells stimulated with plate-bound PD-L1-Fc, CD80-Fc, and α-CD3, did not demonstrate any clear differences in the amount of phosphorylated SHP2 Y542 (pY542), but did have an increase in phosphorylated SHP2 Y580 (pY580) **(Figure 3D)**. Interestingly, cells that were pre-treated with individual blockades of α-PD-1 or α-CTLA-4 did not yield any differences in the amount of pY580 observed but, when both PD-1 and CTLA-4 were blocked, the amount of pY580 was significantly reduced **(Figure 3D, Supplemental Figure 2B)**. Together, these data indicate that PD-1 and CTLA-4 simultaneously contribute to the phosphorylation of TCR-suppressive Y580 that is independent of changes to the Y580-disabling Y542 residue.

Another approach to depict how these treatments impacted the T_reg_ cell populations was to utilize Uniform Manifold Approximation and Projection (UMAP) analysis of concatenated T_reg_ cells from each of the treated groups, excluding the expression of PD-L1 and CTLA-4 from analytical algorithms **(Supplemental Figure 3A)**. Following UMAP analysis, the samples were then unmixed into respective treatment groups and changes in distribution density within the UMAP analysis depicted across the different treatment groups **(Figure 4A)**. Then using the original concatenated UMAP, geometric median fluorescence expression heatmaps were created to show comparative expression of T_reg_-associated proteins (Foxp3, Helios, CD25, and CD122) **(Figure 4B)**, proliferation-associated proteins (cMyc and Ki67) **(Figure 4C)**, activation-associated proteins (Nur77, CD69, CD73) **(Figure 4D)**, and co-stimulation associated proteins **(Figure 4E)**. Compared to isotype treated hosts, the PD-L1 and CTLA-4 blockade treated hosts had increased enrichment in _reg_ions that overlap with Foxp3 and Helios, yet no clear enrichment over the CD25^hi^ _reg_ions of the UMAP. Combination blockade hosts had even further enrichment over the Foxp3^hi^ and Helios^+^ _reg_ions and a comparative reduction of CD25^hi^ cells. **(Figure 4A-B)**. Likewise, either blockade resulted in enrichment in _reg_ions of the UMAP associated with proliferation **(Figure 4C)** or activation **(Figure 4D)**, or expression of B7-family co-stimulation proteins **(Figure 4E)**, with the greatest enrichments occurring in the cohort treated with the combination blockade.

**Figure 4:**
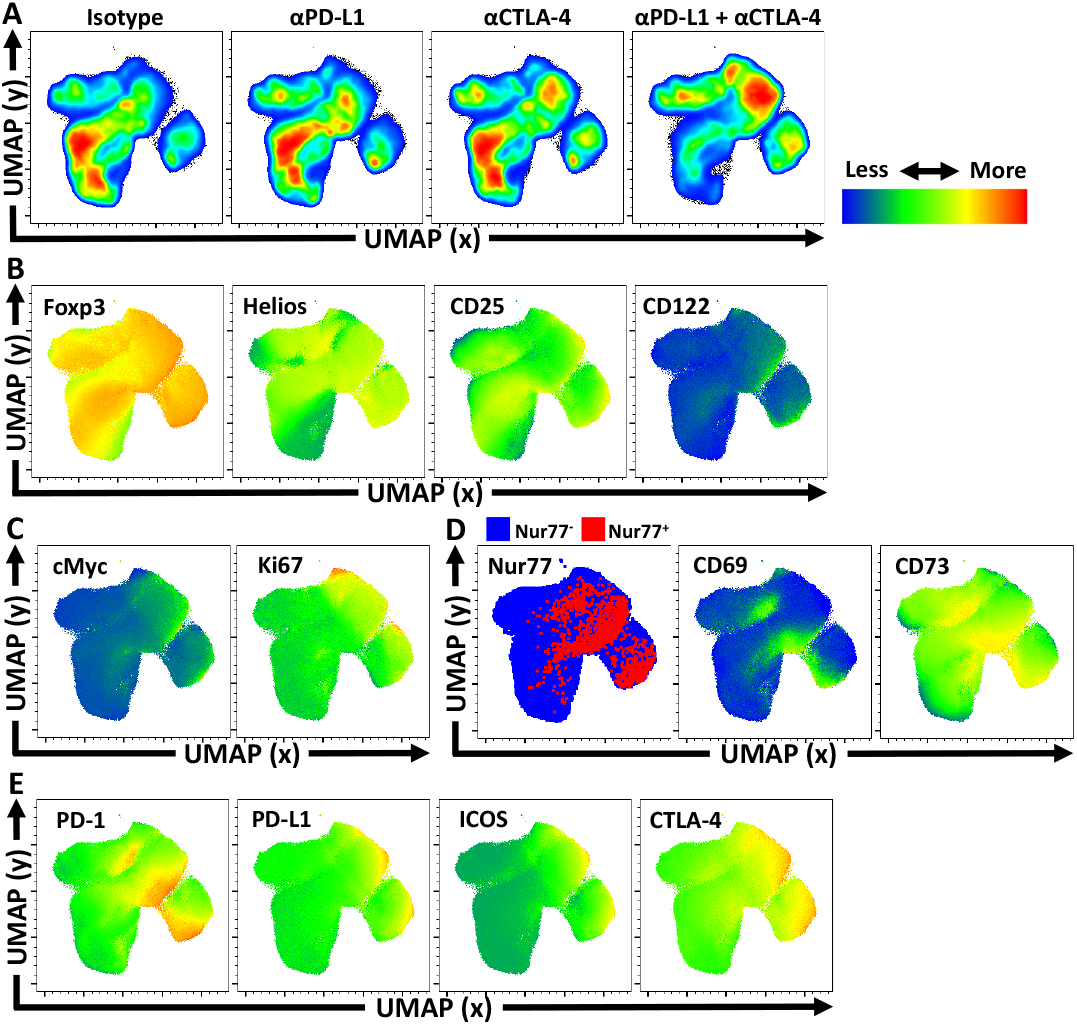
Activated and proliferative T_reg_ compartment phenotypic-shifts following checkpoint blockade. Cohorts of 8 week-old male C57BL/6 mice were given a single intraperitoneal injection of either αPD-L1, or αCTLA-4, or combination αPD-L1 and αCTLA-4, or Isotype control antibody. Splenocytes were harvested for analysis 72 hours later and analyzed via high parameter flow-cytometry. **(A)** Bulk T_reg_ sample data from each treatment group was concatenated and assessed using Uniform Manifold Approximation and Projection (UMAP) analysis **(Supplemental Figure 3A for description)** to produce 2-dimensional plots containing the measured parameters excluding PD-L1 and CTLA-4 from analysis to portray qualitative trends that emerged following treatment. **(A)** Treatment-specific UMAP sub-plots from the concatenated analysis depicting pseudo-color density distribution of T_reg_ cells amongst the UMAP. **(B-D)** Median heatmap expression of individual proteins based on the total concatenated UMAP analysis (including T_reg_ cells from all 4 treatment groups), depicting regions associated with expression the protein label in each box. Heatmap data can then be compared to **4A** to interpret trends based on blockade treatment. **(B)** Expression trends of T_reg_-associated proteins Foxp3, Helios, CD25, and CD122 amongst the T_reg_ compartment. **(C)** Heatmap expression of proliferation associated proteins cMyc and Ki67, with extensive overlap with enriched regions following individual or combination blockade, with more activated and proliferative cells accumulating in the upper right _reg_ion of the UMAP plots, and more quiescent cells in the bottom left _reg_ion of the UMAP plots. **(D)** Nur77^+^ cells (red) overlaid Nur77^-^ cells (blue) amongst the reference UMAP, in addition to heatmaps of activation-associated proteins CD69, and CD73 following blockade treatments. **(E)** Heatmap expression of B7-family costimulatory proteins, PD-1, PD-L1, ICOS, and CTLA-4, with enrichment in the _reg_ions correlating to blockade treatment.

One of the major effects of blockade treatment was on the number of T_reg_ cells that expressed Ki67 and cMyc with the most prominent effect observed when both PD-L1 and CTLA-4 were blocked **(Figure 5A)**. This observation was consistent with the increased number of the PD-1^+^ T_reg_ subsets (PD-1^low^, PD-1^hi^) **(Figure 5B)**. Additionally, TCR stimulation of T_reg_ cells is associated with maintenance of Foxp3 expression^12^, and antagonism of TCR activity by treatment of mice for 4 days with the tacrolimus (FK506)^42^ resulted in a reduced MFI of Foxp3 in T_reg_ cells **(Figure 5C)**. In contrast, the blockade of PD-L1 or CTLA-4 resulted in an overall increase in the MFI of Foxp3 amongst the bulk T_reg_ compartment, with the combination blockade having the greatest enhancement **(Figure 5D, Supplemental Figure 3B for individual eT**_**reg**_ **blockade comparisons)**. Combined with the numerical, phenotypic and phos-data sets these results highlight that PD-1 and CTLA-4 additively contribute restrict the population of TCR-driven effector T_reg_ cells.

**Figure 5:**
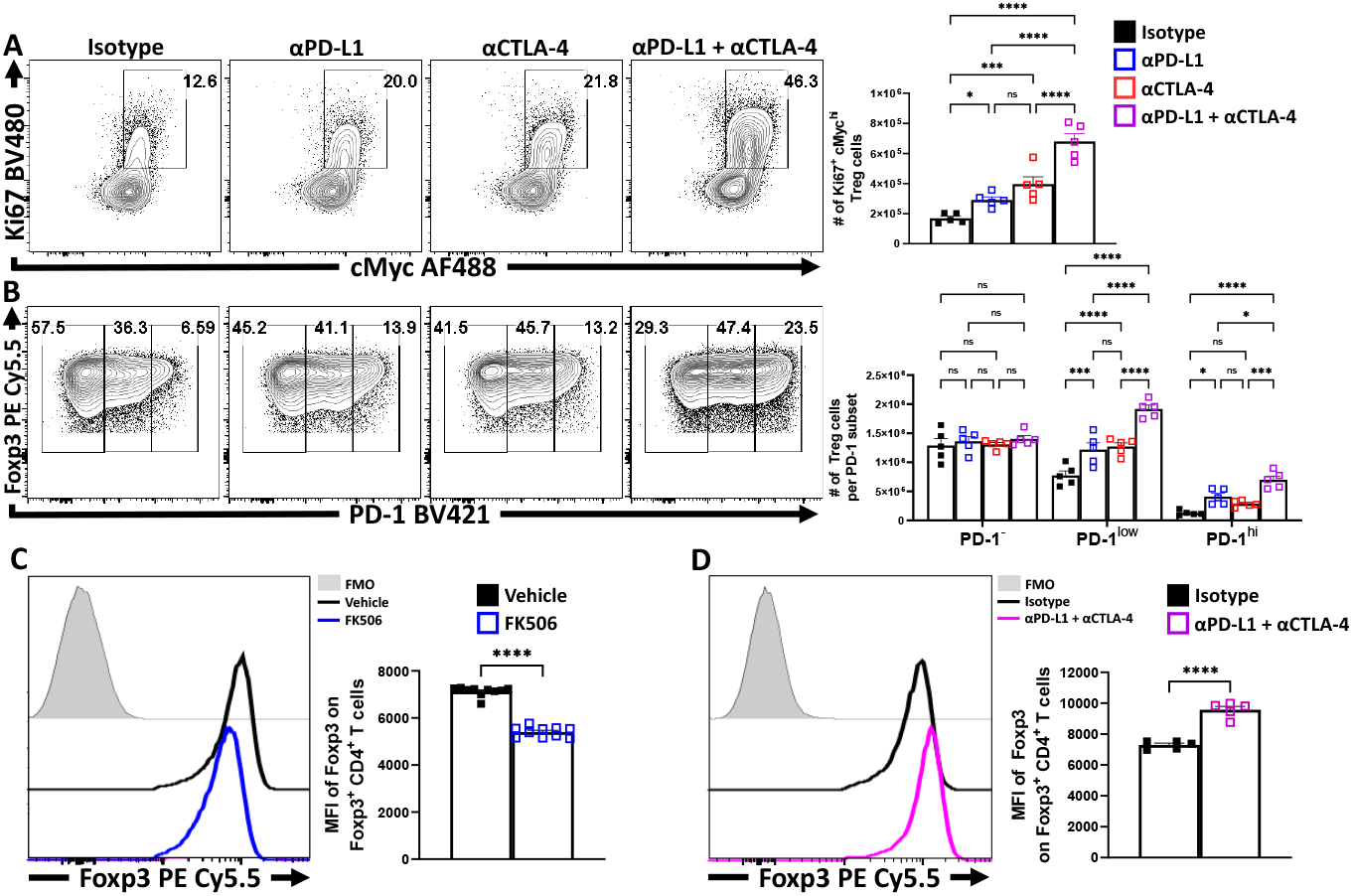
Blockade of PD-L1 and CTLA-4 additively drive enrichment and proliferation of the eT_reg_ compartment. Cohorts of 8 week-old male C57BL/6 mice were given a single intraperitoneal injection of either αPD-L1, or αCTLA-4, or combination αPD-L1 and αCTLA-4, or Isotype control antibody. Splenocytes were harvested for analysis 72 hours later and analyzed via high parameter flow-cytometry. **(A)** Flow plots comparing the proportion and number of Ki67^+^ cMyc^hi^ T_reg_ cells following individual or combination PD-L1/CTLA-4 blockade treatment *(n = 5/group, 1-way ANOVA with Fisher’s LSD individual comparisons test, * = p < 0*.*05, *** = p < 0*.*001, **** = p < 0*.*0001, 3 experimental replicates)*. **(B)** Flow plots comparing the proportions and number of PD-1^-^, PD-1^low^, and PD-1^hi^ T_reg_ cells following individual or combination blockade treatment, with enrichments occurring within the PD-1^+^ eT_reg_ associated subsets following blockade, the greatest of which occur with combination blockade treatment (*2-way ANOVA with Sidak’s multiple comparisons test, * = p < 0*.*05, *** = p < 0*.*001, **** = p < 0*.*0001, 3 experimental replicates)*. **(C)** Cohorts of 8 week-old male C57BL/6 mice were treated once daily for 4 days with subcutaneous injections of PBS/vehicle or Tacrolimus (FK506), and splenocytes were harvested and analyzed via flow cytometry. Comparative histograms of T_reg_ cells from vehicle control and FK506 treated mice demonstrating decreases in the gMFI of Foxp3 in T_reg_ cells in FK506 treated hosts *(n = 10/group two-tailed unpaired student’s t-test, **** = p < 0*.*0001, 2 experimental replicates)*. **(D)** Comparative histograms of T_reg_ cells from isotype and αPD-L1/αCTLA-4 combination blockade treated mice demonstrating increases in the gMFI of Foxp3 in T_reg_ cells from blockade treated hosts *(n = 5/group two-tailed unpaired student’s t-test, **** = p < 0*.*0001, 3 experimental replicates). All data presented are means +/- SEM and show individual data points*.

Finally, to determine the impact of PD-1 and CTLA-4 on T_reg_ function splenocytes from treated hosts were stimulated and the ability to produce IL-10 assessed. Either PD-L1 blockade or CTLA-4 blockade resulted in an increase in the number of IL-10^+^ T_reg_ cells, with the combination blockade resulting in the greatest increase **(Figure 6A)**. IL-10 is an inhibitor of the ability of macrophages and DC to express CD80 and MHC class II whereas the ability of CTLA-4 to bind to and strip CD80 from these cells can reduce costimulation^32^. Evaluation of these compartments ex vivo **(Supplemental Figure 4)** showed that cDC2s and macrophages had varied expression of MHC class II and CD80. The blockade of CTLA-4 or PD-L1 alone did not alter DC expression of class II, while the combined blockade had modest but consistent trends of decreasing MHC class II expression in cDC2s and macrophages **(Figure 6B)**. In contrast, PD-L1 blockade alone reduced CD80 on cDC2s and macrophages, yet consistent with the ability of CTLA-4 to strip CD80, CTLA-4 blockade resulted in increased CD80 expression **(Figure 6C)**. When blockade treatments were combined, the effects of anti-PD-L1 were dominant with reduction in the expression of CD80. This result established that not only do these treatments alter the T_reg_ compartment, but this correlates with alterations of other cell types that are known to be impacted by eT_reg_ cells.

**Figure 6:**
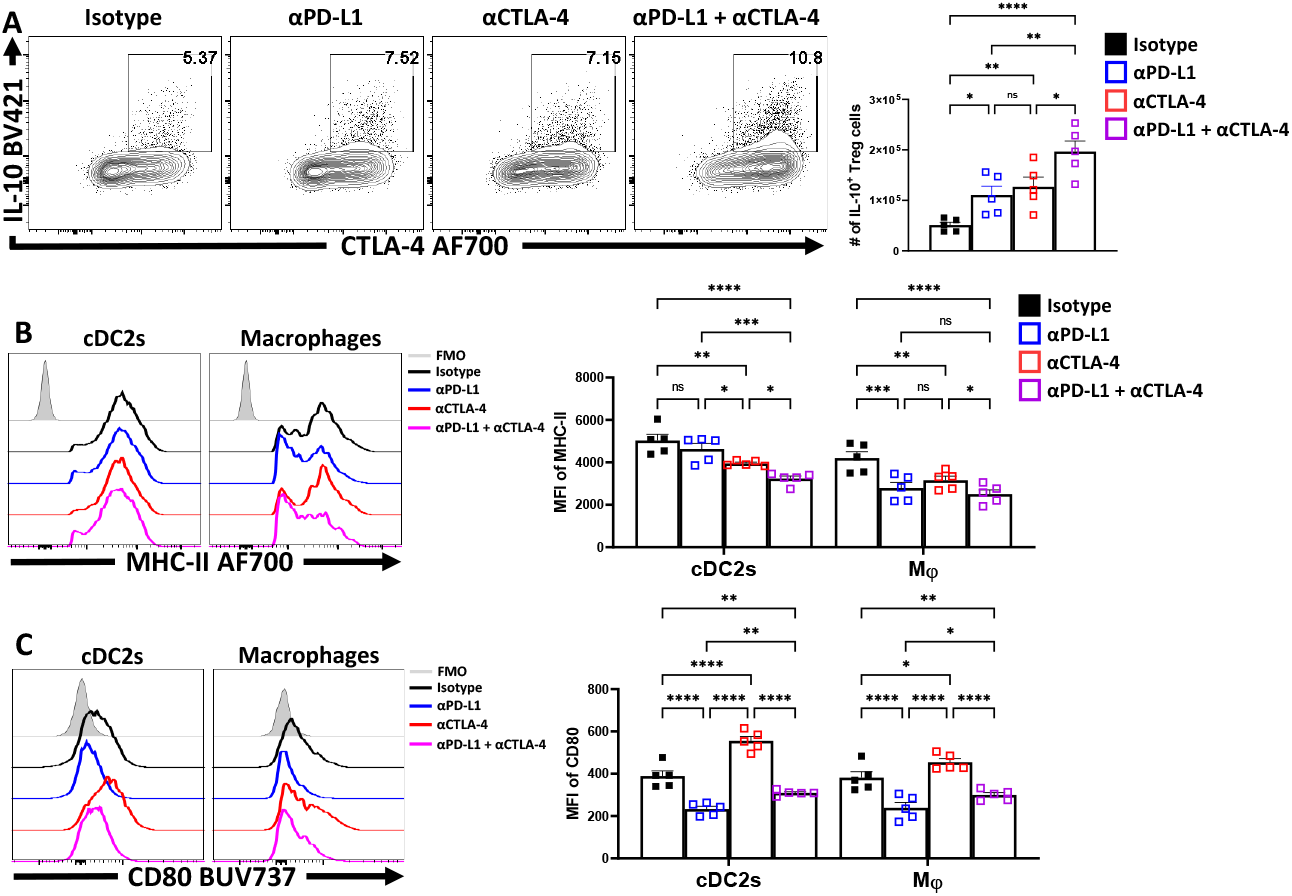
Combo-blockade of PD-L1 and CTLA-4 drives a myeloid-suppressive T_reg_ environment. 8 week-old male C57BL/6 mice were given a single intraperitoneal injection of either αPD-L1, or αCTLA-4, or combination αPD-L1 and αCTLA-4, or Isotype control antibody. At 72 hours following treatment, their splenocytes were harvested, and stimulated with PMA/Ionomycin for cytokine staining and analyzed via flow cytometry. **(A)** Plots depicting the expression of IL-10 and CTLA-4 on bulk T_reg_ cells from each treatment group, with increases in IL-10^+^ CTLA-4^hi^ T_reg_ cells from single and combo blockade treated hosts *(n = 5/group, 1-way ANOVA with Fisher’s LSD individual comparisons test, * = p < 0*.*05, ** = p < 0*.*01, **** = p < 0*.*0001, 2 experimental replicates)*. **(B)** Plots comparing MHC-II expression on cDC2s and Macrophages **(Supplemental Figure 4 for description)** following blockade treatments, with decreasing trends MHC-II with combo blockade *(n = 5/group, 2-way ANOVA with Fisher’s LSD individual comparisons test, * = p < 0*.*05, *** = p < 0*.*001, **** = p < 0*.*0001, 3 experimental replicates)* **(C)** Ex-vivo staining of splenocytes evaluating the expression of CD80 on cDC2s (CD3^-^, B220^-^, CD19^-^, NK1.1^-^, Ly6G^-^, CD64^-^, CD11c^+^, MHC-II^+^, SIRPα^+^), and macrophages (CD3^-^, B220^-^, CD19^-^, NK1.1^-^, Ly6G-, CD64^+^, CD11b^+^, MHC-II^+^, Ly6C^low^), demonstrating changes to surface CD80 based on blockade treatment *(n = 5/group, 2-way ANOVA with Fisher’s LSD individual comparisons test, * = p < 0*.*05, ** = p < 0*.*01, **** = p < 0*.*0001, 3 experimental replicates). All data presented are means +/- SEM and show individual data points*.

## Discussion

The focus on the role of PD-1 and CTLA-4 in limiting effector T cell responses has revealed that the expression of these molecules is a byproduct of repeated TCR stimulation over time^36,43,44^. In this context, it appears that similar rules apply to T_reg_ cells which receive and require ongoing TCR signaling to maintain the expression of Foxp3 and their suppressive capacity^12^. Thus, mitigating PD-1 and CTLA-4 signaling results in greater overall expression of Foxp3 while even short-term blockade of TCR activation reduced Foxp3 expression. Earlier models suggested that PD-1 and CTLA-4 are associated with the suppressive functions of T_reg_ cells ^45,46^, but a consensus is emerging that these inhibitory receptors can individually restrict T_reg_ capacity and suppressive function during autoimmune disease^21,27^, cancer^22^ and infection ^28^. The studies presented here focus on the impact of these potentially-overlapping pathways on T_reg_ cell homeostasis and in particular on the differences between eT_reg_ and cT_reg_ cell populations. We now appreciate that while the cT_reg_ compartment makes greater use of STAT5 signaling cytokines such as IL-2 for maintenance^5,14,47^, the eT_reg_ compartment is more dependent on TCR-mediated activation and co-stimulation to survive. Indeed, we present data which indicates that the eT_reg_ subset has higher basal signals of pZAP70, pAKT, and pmTOR, than the cT_reg_ compartment. However, eT_reg_ cells also express PD-1 and CTLA-4, related proteins which overlap in their ability to engage SHP2 signaling to antagonize T cell activation^30,48,49^. This observation implies that the eT_reg_ subset has an enhanced capacity to respond to TCR signals while being sensitive to negative signals from PD-1 or CTLA-4. The relevance of tonic signals through these IRs was shown by the finding that mitigation of both PD-1 and CTLA-4 signaling pathways reduces suppressive pSHP2 Y580, which coincided with increased Foxp3 expression and numbers of eT_reg_ cells and enhanced IL-10 production.

While PD-1 and CTLA-4 are related B7 family members, engage SHP2 signaling, and seem to additively limit eT_reg_ proliferation and function, there is still distinction to their suppressive mechanisms. Thus, PD-1 accumulates on the cell surface and is accessible to PD-L1 ligation^50^. and PD-1 appears to act in cis to limit T cell activation. For T_reg_ cells, blockade of this pathway resulted in enhanced numbers and IL-10 production and was associated with reduced accessory cell expression of CD80 and MHC class II. In contrast, the majority of CTLA-4 is stored intracellularly and is translocated to the surface upon TCR stimulation^51^ where it can provide negative costimulatory signals^52^. In addition, the ability of CTLA-4 to bind with high affinity to CD80 means that it can outcompete the ability of CD28 to provide costimulation and can actively restrict APC function through CTLA-4 mediated trogocytosis of CD80^32^. Thus, CTLA-4 is an invoked off switch which can simultaneously act in cis and trans to limit eT_reg_ cells. Interestingly, this complex biology is apparent in the studies presented here: α-PD-L1 treatment alone drove a reduction in CD80 expression by cDC2s and macrophages, while α-CTLA-4 treatment still drove an enrichment of T_reg_ cells yet resulted in a significant increase in myeloid expression of CD80 (consistent with reduced trogocytosis). Nevertheless, that combination blockade of PD-L1 and CTLA-4 resulted in a reduction of myeloid CD80 expression, which suggest that the increased number of T_reg_ cells and their production of IL-10 is sufficient to surpass the effects of CTLA4 on CD80 levels.

The past twenty years has witnessed an increased utilization of immunotherapeutic drugs to enhance immune mediated control of certain cancers or to limit autoimmune inflammation. The blockade of PD-1 or CTLA-4 or the use of CTLA4-Ig or all examples of clinical interventions to impact effector T cell responses that are directly relevant to the pathways that we show here are also relevant to eT_reg_ cells. However, these treatment strategies do not always prove effective and their impact of T_reg_ cells may in part explain some of this heterogeneity in clinical outcome^22,53,54^. Perhaps, the ability to specifically target these pathways (either to agonize or block) on eT_reg_ cells can be used as an immunotherapeutic strategy to enhance T_reg_ function to treat immunopathological diseases or select against T_reg_ mediated suppression in the context of infection or cancer.

## Acknowledgements

This project was supported by NIAID R01 AI125563 & R01 AI41158, awarded to Christopher Hunter.

Joseph Perry was supported by training grant: T32-CA-009140. Thank you to Keith Burton for your patience and support.

## Author Contributions Statement

J.A.P conceptualized the project, designed/executed all experiments, performed data analysis, figure production, and authored the paper. Z.L, J.T.C, A.P.H, B.B.D, L.S., and K.O., aided in data collection, provided conceptual feedback _reg_arding experimental design, data analysis, and manuscript editing. D.A.C. directly supervised experimental execution, interpretation, and presentation of data. C.A.H. supervised the project in entirety.

The authors of this manuscript have reviewed the work presented here in raw form and approve it. Appropriate statistical tests were utilized, and the figures are correct depictions of the data without manipulation aside from resizing for press. The policies set by the journal _reg_arding materials, data sharing, ethical animal use, and conflicts of interest have been followed. We are confident the conclusions presented here are based on accurate interpretations of the data collected for this study.

## Competing Interests Statement

The authors have declared no competing interests. Subject to Patent disclosure.

---

***Supplemental Figure 1: T***_***reg***_ ***subsetting***.

**(A)** Flow cytometry sub-gating example strategy identifying T_reg_ cells, utilizing splenocytes from an 8 week-old male C57BL/6 mouse. **(B)** Gating strategy to identify PD-1^+^ CTLA-4^hi^ (eT_reg_) vs PD-1^-^ CTLA-4^low^ (cT_reg_) subsets. **(C)** Splenocytes from an 8 week-old and 16 week-old C57BL/6 mice were evaluated for their proportions of PD-1^+^ CTLA-4^hi^ eT_reg_ cells *(two-tailed unpaired student’s t-test, ** = p < 0*.*01). All data presented are means +/- SEM and show individual data points*.

***Supplemental Figure 2: CD4 T cell enrichment and SHP2 phosphorylation results***.

**(A)** Flow plots of from 8 week-old male C57BL/6 mouse splenocytes assessing CD3^+^ CD4^+^ T cell proportions following MACS enrichment *(two-tailed paired student’s t-test, **** = p < 0*.*0001, 2 experimental replicates)*. **(B)** Enriched bulk CD4^+^ T cells were treated with either αPD-1, or αCTLA-4, or combination αPD-1 and αCTLA-4, or Isotype control antibody, and then stimulated with plate-bound α-CD3, PD-L1-Fc, and CD80-Fc and phospho- stained. Depicted are histogram comparisons of the T_reg_ subset (CD4^+^ Foxp3^+^) comparing gMFI of p-SHP2 at tyrosine residues Y542 and Y580 on T_reg_ cells *(n = 5/group, 1-way ANOVA with Fisher’s LSD individual comparisons test, * = p < 0*.*05, ** = p < 0*.*01, *** = p < 0*.*001, **** = p < 0*.*0001, 2 experimental replicates). All data presented are means +/- SEM and show individual data points*.

***Supplemental Figure 3: UMAP and eT***_***reg***_ ***Foxp3 MFI following checkpoint blockade***.

**(A)** Flow cytometry sub-gating example strategy identifying T_reg_ cells, utilizing splenocytes from an 8 week-old male C57BL/6 mouse. The UMAP was then generated using concatenated T_reg_ cells from 8 week-old male C57BL/6 mice that were given a single intraperitoneal injection of either αPD-L1, or αCTLA- 4, or combination αPD-L1 and αCTLA-4, or Isotype control antibody. The calculated fluorescence factors that generated the UMAP are depicted. **(B)** Comparative histograms depicting the gMFI of Foxp3 on PD-1^+^ CTLA-4^hi^ eT_reg_ cells following checkpoint blockade *(n = 5/group, 1-way ANOVA with Fisher’s LSD individual comparisons test, * = p < 0*.*05, ** = p < 0*.*01, **** = p < 0*.*0001, 2 experimental replicates). All data presented are means +/- SEM and show individual data points*.

***Supplemental Figure 4: Myeloid Gating***.

**(A)** Splenocytes from 8 week-old male C57BL/6 mice were analyzed via flow cytometry across multiple leukocyte populations as depicted: B cells (CD3^-^, B220^+^, CD19^+^, MHC-II^+^), cDC1s (CD3^-^, B220^-^, CD19^-^, NK1.1^-^, Ly6G^-^, CD64^-^, CD11c^+^, MHC-II^+^, XCR1^+^), cDC2s (CD3^-^, B220^-^, CD19^-^, NK1.1^-^, Ly6G^-^, CD64^-^, CD11c^+^, MHC-II^+^, SIRPα^+^), and macrophages (CD3^-^, B220^-^, CD19^-^, NK1.1^-^, Ly6G^-^, CD64^+^, CD11b^+^, MHC-II^+^, Ly6C^low^).

## References

1. Fontenot, J. D., Gavin, M. A. & Rudensky, A. Y. Foxp3 programs the development and function of CD4+CD25+ _reg_ulatory T cells. Nat. Immunol. 4, 330–336 (2003).

2. Josefowicz, S. Z., Lu, L.-F. & Rudensky, A. Y. Regulatory T Cells: Mechanisms of Differentiation and Function. Annu. Rev. Immunol. 30, 531–564 (2012).

3. Sakaguchi, S. et al. Regulatory T Cells and Human Disease. Annu. Rev. Immunol. 38, 541–566 (2020).

4. Sakaguchi, S., Wing, K. & Miyara, M. Regulatory T cells - A brief history and perspective. Eur. J. Immunol. 37, 116–123 (2007).

5. Smigiel, K. S. et al. CCR7 provides localized access to IL-2 and defines homeostatically distinct _reg_ulatory T cell subsets. J. Exp. Med. 211, 121–136 (2014).

6. Liston, A. & Gray, D. H. D. Homeostatic control of _reg_ulatory T cell diversity. Nat. Rev. Immunol. 14, 154–165 (2014).

7. Bilate, A. M. & Lafaille, J. J. Induced CD4 + Foxp3 + Regulatory T Cells in Immune Tolerance. Annu. Rev. Immunol. 30, 733–758 (2012).

8. Fan, M. Y. et al. Differential Roles of IL-2 Signaling in Developing versus Mature Tregs. Cell Rep. 25, 1204-1213.e4 (2018).

9. Kieback, E. et al. Thymus-Derived Regulatory T Cells Are Positively Selected on Natural Self-Antigen through Cognate Interactions of High Functional Avidity. Immunity 44, 1114–1126 (2016).

10. Bautista, J. L. et al. Intraclonal competition limits the fate determination of _reg_ulatory T cells in the thymus. Nat. Immunol. 10, 610–617 (2009).

11. Owen, D. L., Sjaastad, L. E. & Farrar, M. A. Regulatory T Cell Development in the Thymus. J. Immunol. 203, 2031–2041 (2019).

12. Levine, A. G., Arvey, A., Jin, W. & Rudensky, A. Y. Continuous requirement for the TCR in _reg_ulatory T cell function. Nat. Immunol. 15, 1070–1078 (2014).

13. Wakamatsu, E., Mathis, D. & Benoist, C. Convergent and divergent effects of costimulatory molecules in conventional and _reg_ulatory CD4+ T cells. Proc. Natl. Acad. Sci. U. S. A. 110, 1023–1028 (2013).

14. Kornete, M., Mason, E., Istomine, R. & Piccirillo, C. A. KLRG1 expression identifies short-lived Foxp3(+) T_reg_ effector cells with functional plasticity in islets of NOD mice. Autoimmunity 50, 1–9 (2017).

15. Wildin, R. S. et al. X-linked neonatal diabetes mellitus, enteropathy and endocrinopathy syndrome is the human equivalent of mouse scurfy. Nat. Genet. 27, 18–20 (2001).

16. Bennett, C. L. et al. The immune dys_reg_ulation, polyendocrinopathy, enteropathy, X-linked syndrome (IPEX) is caused by mutations of FOXP3. Nat. Genet. 27, 20–21 (2001).

17. Wilson, E. H., Wille-Reece, U., Dzierszinski, F. & Hunter, C. A. A critical role for IL-10 in limiting inflammation during toxoplasmic encephalitis. J. Neuroimmunol. 165, 63–74 (2005).

18. Warunek, J. et al. Tbet Expression by Regulatory T Cells Is Needed to Protect against Th1-Mediated Immunopathology during Toxoplasma Infection in Mice. ImmunoHorizons 5, 931–943 (2021).

19. Husebye, E. S., Anderson, M. S. & Kämpe, O. Autoimmune Polyendocrine Syndromes. N. Engl. J. Med. 378, 1132–1141 (2018).

20. Benson, A. et al. Microbial Infection-Induced Expansion of Effector T Cells Overcomes the Suppressive Effects of Regulatory T Cells via an IL-2 Deprivation Mechanism. J. Immunol. 188, 800–810 (2012).

21. Paterson, A. M. et al. Deletion of CTLA-4 on _reg_ulatory T cells during adulthood leads to resistance to autoimmunity. J. Exp. Med. 212, 1603–1621 (2015).

22. Kamada, T. et al. PD-1 + _reg_ulatory T cells amplified by PD-1 blockade promote hyperprogression of cancer. Proc. Natl. Acad. Sci. 116, 201822001 (2019).

23. Xu, T., Lu, J. & An, H. The relative change in _reg_ulatory T cells / T helper lymphocytes ratio as parameter for prediction of therapy efficacy in metastatic colorectal cancer patients. Oncotarget 8, 109079–109093 (2017).

24. Wang, B. et al. Combination cancer immunotherapy targeting PD-1 and GITR can rescue CD8+ T cell dysfunction and maintain memory phenotype. Sci. Immunol. 3, 1–14 (2018).

25. Oldenhove, G. et al. Decrease of Foxp3+ T_reg_ Cell Number and Acquisition of Effector Cell Phenotype during Lethal Infection. Immunity 31, 772–786 (2009).

26. Hernandez, R., Põder, J., LaPorte, K. M. & Malek, T. R. Engineering IL-2 for immunotherapy of autoimmunity and cancer. Nat. Rev. Immunol. 0123456789, (2022).

27. Tan, C. L. et al. PD-1 restraint of _reg_ulatory T cell suppressive activity is critical for immune tolerance. J. Exp. Med. 218, (2020).

28. Perry, J. A. et al. PD-L1–PD-1 interactions limit effector _reg_ulatory T cell populations at homeostasis and during infection. Nat. Immunol. 23, 743–756 (2022).

29. Simpson, T. R. et al. Fc-dependent depletion of tumor-infiltrating _reg_ulatory t cells co-defines the efficacy of anti-CTLA-4 therapy against melanoma. J. Exp. Med. 210, 1695–1710 (2013).

30. Schneider, H. & Rudd, C. E. Tyrosine phosphatase SHP-2 binding to CTLA-4: Absence of direct YVKM/YFIP motif recognition. Biochem. Biophys. Res. Commun. 269, 279–283 (2000).

31. Hui, E. et al. T cell costimulatory receptor CD28 is a primary target for PD-1–mediated inhibition. Science (80-.). 355, 1428–1433 (2017).

32. Tekguc, M., Wing, J. B., Osaki, M., Long, J. & Sakaguchi, S. T_reg_-expressed CTLA-4 depletes CD80/CD86 by trogocytosis, releasing free PD-L1 on antigen-presenting cells. Proc. Natl. Acad. Sci. 118, (2021).

33. Hünig, T., Beyersdorf, N. & Kerkau, T. CD28 co-stimulation in T-cell homeostasis: a recent perspective. ImmunoTargets Ther. 111 (2015) doi:10.2147/ITT.S61647.

34. Pen, J. J. et al. Interference with PD-L1/PD-1 co-stimulation during antigen presentation enhances the multifunctionality of antigen-specific T cells. Br. Dent. J. 217, 262–271 (2014).

35. Li, C. PD-1 and CTLA-4 Mediated Inhibitory Signaling for T cell Exhaustion during Chronic Viral Infections. J. Clin. Cell. Immunol. 01, (2013).

36. Wherry, E. J. & Kurachi, M. Molecular and cellular insights into T cell exhaustion. Nat. Rev. Immunol. 15, 486–499 (2015).

37. Samusik, N., Good, Z., Spitzer, M. H., Davis, K. L. & Nolan, G. P. Automated mapping of phenotype space with single-cell data. Nat. Methods 13, 493–496 (2016).

38. Schmidt, E. V. The role of c-myc in cellular growth control. Oncogene 18, 2988–2996 (1999).

39. Dose, M. et al. c-Myc mediates pre-TCR-induced proliferation but not developmental progression. Blood 108, 2669–2677 (2006).

40. Hui, E. et al. T cell costimulatory receptor CD28 is a primary target for PD-1–mediated inhibition. Science (80-.). 4, eaaf1292 (2017).

41. Lu, W., Gong, D., Bar-Sagi, D. & Cole, P. A. Site-specific incorporation of a phosphotyrosine mimetic reveals a role for tyrosine phosphorylation of SHP-2 in cell signaling. Mol. Cell 8, 759–769 (2001).

42. Ho, S. et al. The mechanism of action of cyclosporin A and FK506. Clin. Immunol. Immunopathol. 80, 1433–1439 (1996).

43. Chemnitz, J. M., Parry, R. V., Nichols, K. E., June, C. H. & Riley, J. L. SHP-1 and SHP-2 Associate with Immunoreceptor Tyrosine-Based Switch Motif of Programmed Death 1 upon Primary Human T Cell Stimulation, but Only Receptor Ligation Prevents T Cell Activation. J. Immunol. 173, 945–954 (2004).

44. Blackburn, S. D. et al. Co_reg_ulation of CD8+ T cell exhaustion by multiple inhibitory receptors during chronic viral infection. Nat. Immunol. 10, 29–37 (2009).

45. Keir, M. E., Butte, M. J., Freeman, G. J. & Sharpe, A. H. PD-1 and Its Ligands in Tolerance and Immunity. Annu. Rev. Immunol. 26, 677–704 (2008).

46. Wing, K. et al. CTLA-4 control over Foxp3+ _reg_ulatory T cell function. Science (80-.). 322, 271–275 (2008).

47. Sprouse, M. L. et al. Cutting Edge: Low-Affinity TCRs Support Regulatory T Cell Function in Autoimmunity. J. Immunol. 200, 909–914 (2018).

48. Walunas, T. L. et al. CTLA-4 can function as a negative _reg_ulator of T cell activation. Immunity 1, 405–413 (1994).

49. Walker, L. S. K. PD-1 and CTLA-4: Two checkpoints, one pathway? Sci. Immunol. 2, 1–5 (2017).

50. Horne-Debets, J. M. et al. PD-1 dependent exhaustion of CD8+T cells drives chronic malaria. Cell Rep. 5, 1204–1213 (2013).

51. Schneider, H. & Rudd, C. E. Diverse mechanisms _reg_ulate the surface expression of immunotherapeutic target CTLA-4. Front. Immunol. 5, 1–10 (2014).

52. Guntermann, C. & Alexander, D. R. CTLA-4 Suppresses Proximal TCR Signaling in Resting Human CD4 + T Cells by Inhibiting ZAP-70 Tyr 319 Phosphorylation: A Potential Role for Tyrosine Phosphatases. J. Immunol. 168, 4420–4429 (2002).

53. Marin-Acevedo, J. A., Kimbrough, E. M. O. & Lou, Y. Next generation of immune checkpoint inhibitors and beyond. J. Hematol. Oncol. 14, 1–29 (2021).

54. Fares, C. M., Van Allen, E. M., Drake, C. G., Allison, J. P. & Hu-Lieskovan, S. Mechanisms of Resistance to Immune Checkpoint Blockade: Why Does Checkpoint Inhibitor Immunotherapy Not Work for All Patients? Am. Soc. Clin. Oncol. Educ. B. 147–164 (2019) doi:10.1200/edbk_240837.

