## Supplementary figures and images for "PD-1 and CTLA-4 exert additive control of effector regulatory T cells"

### Supplemental Figures 1-4

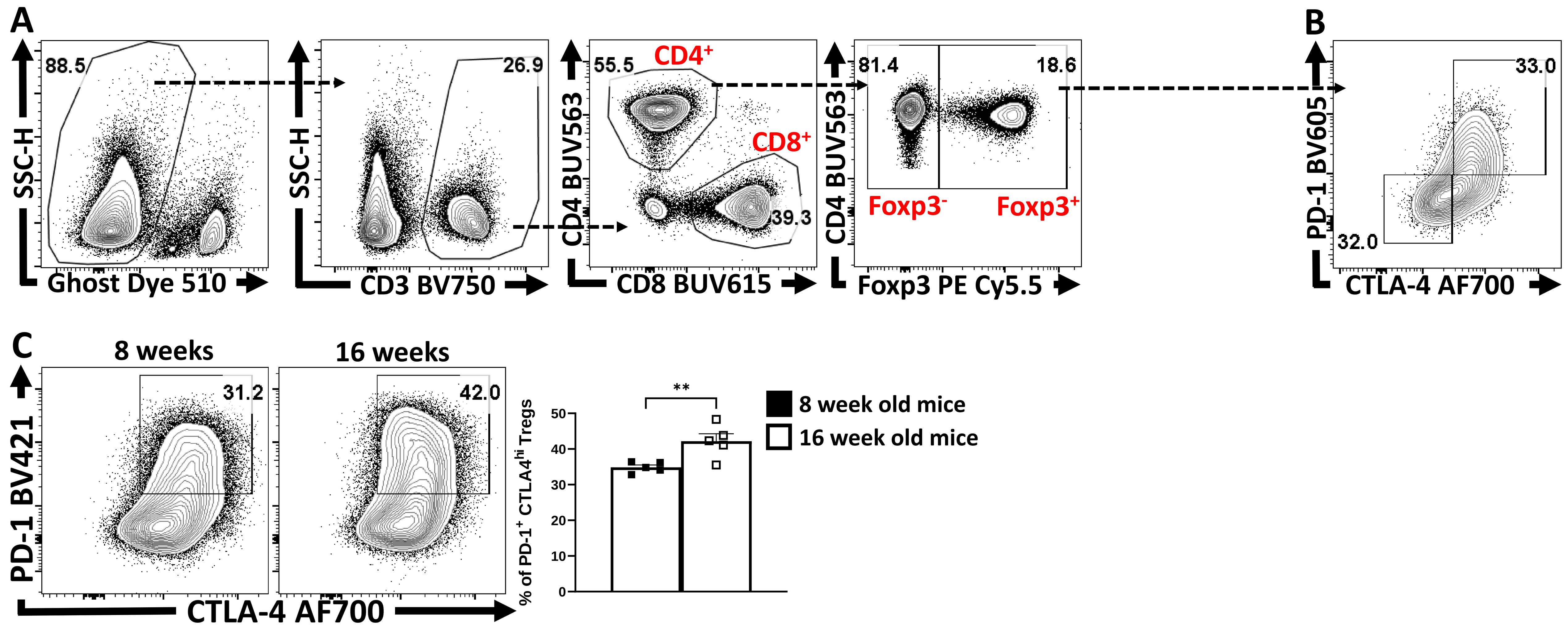

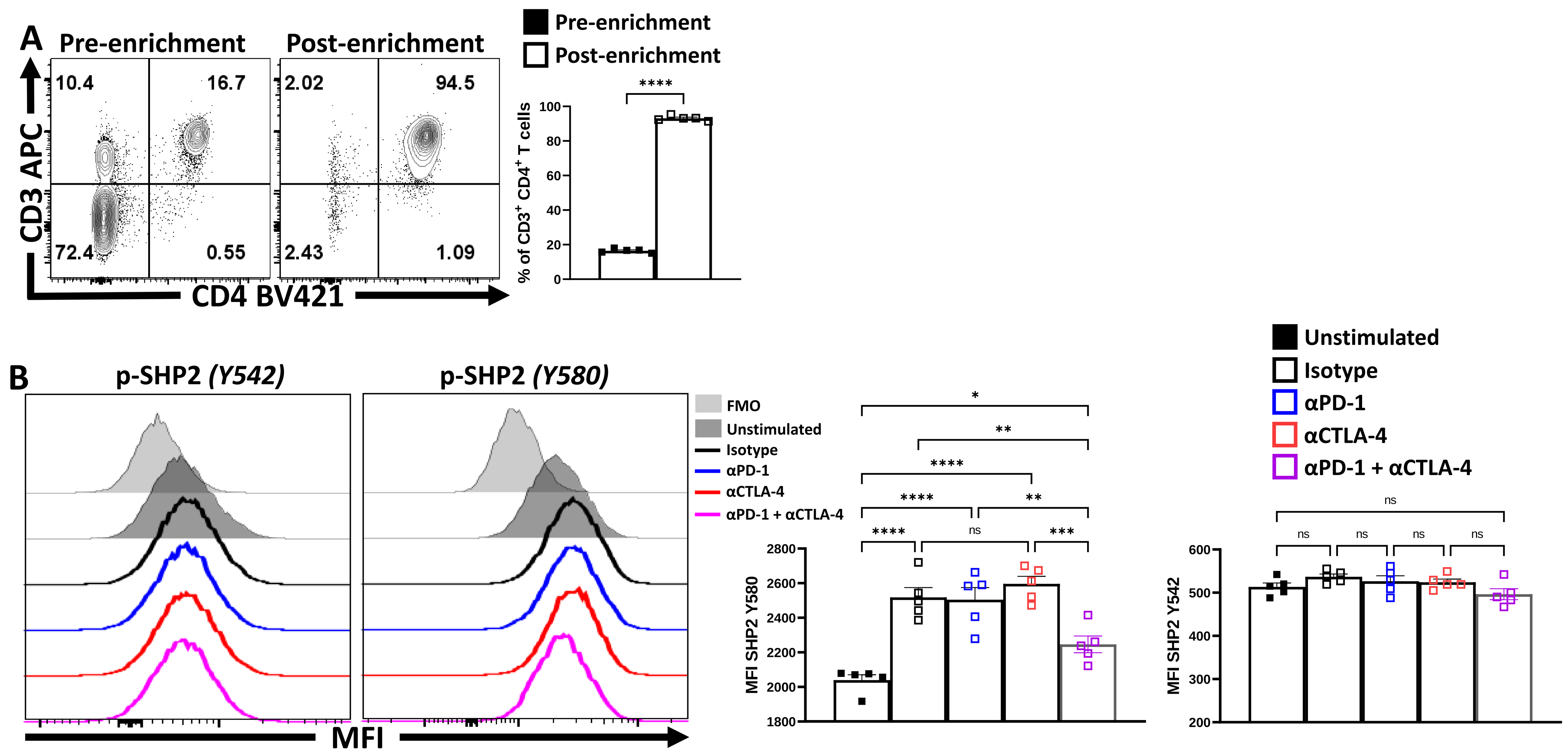

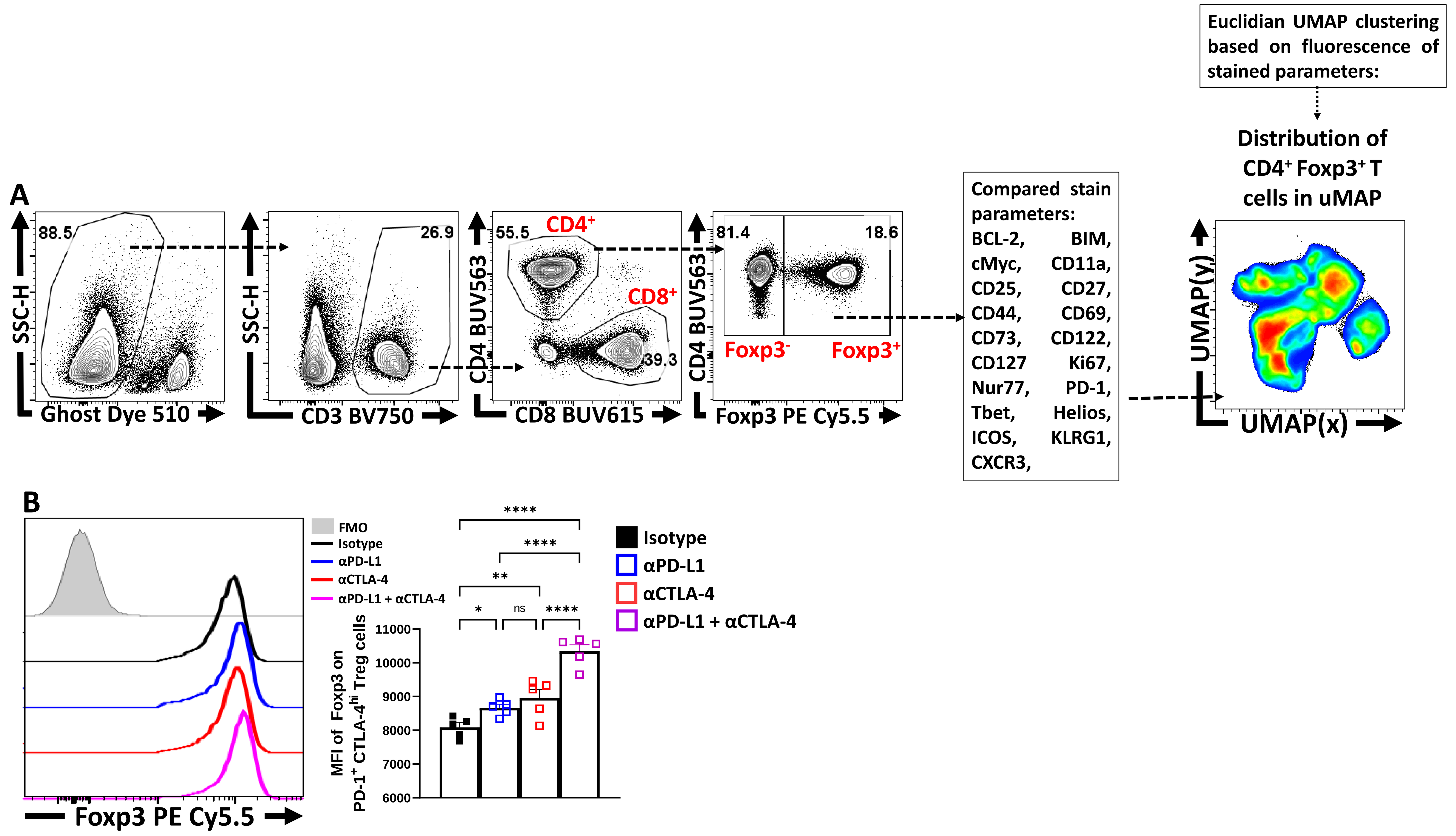

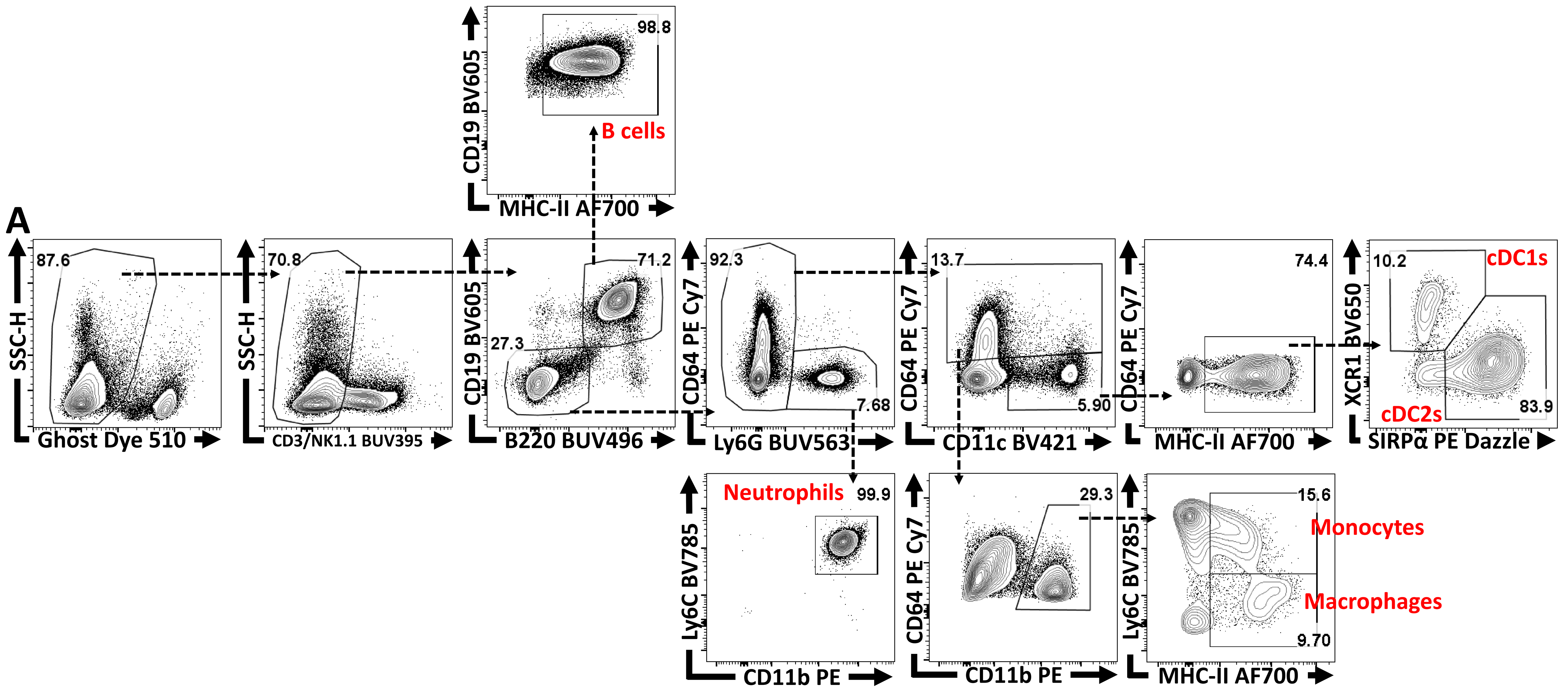

### Supplemental Table 1

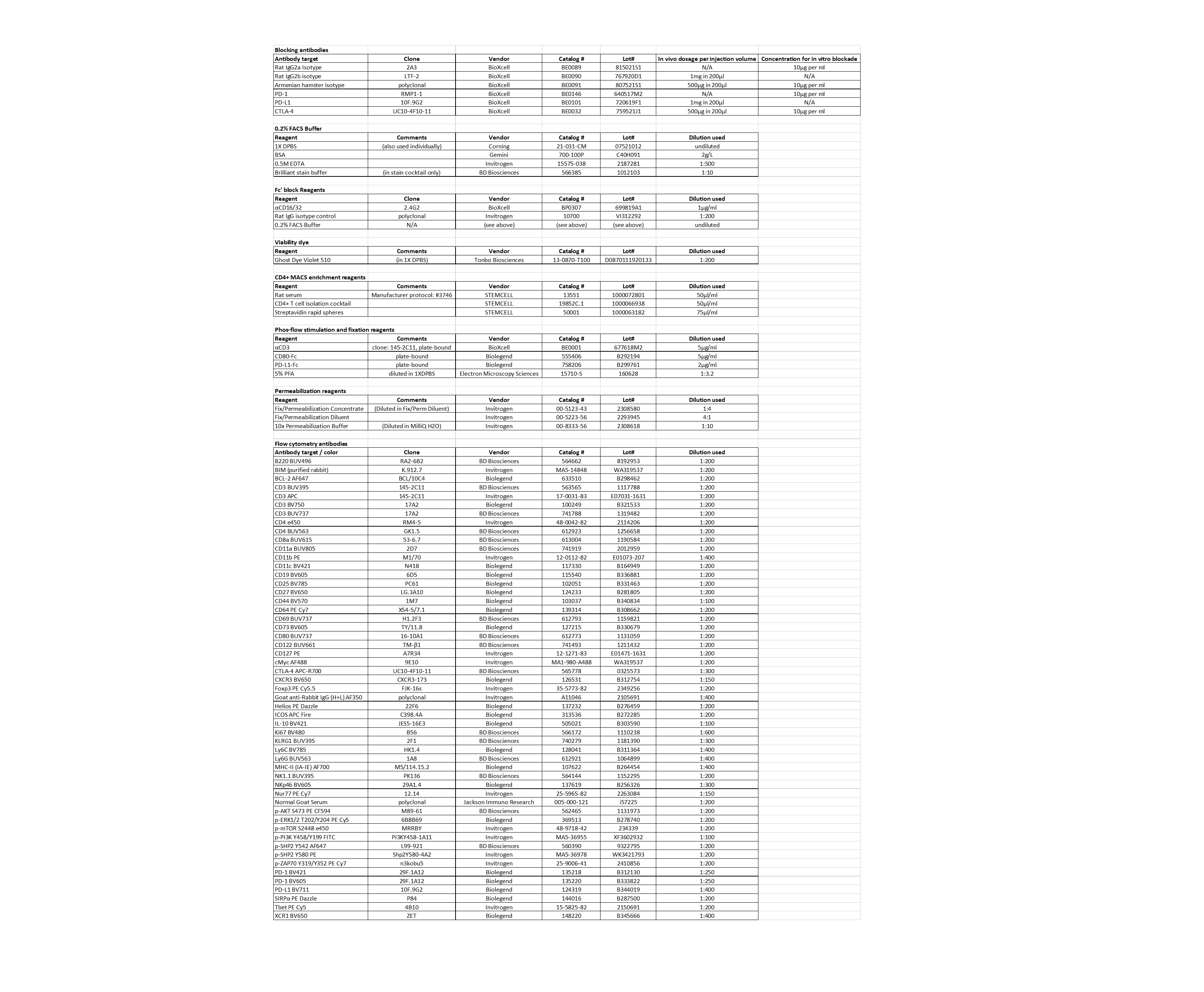
